# Criticality of plasma membrane lipids reflects activation state of macrophage cells

**DOI:** 10.1101/680157

**Authors:** Eugenia Cammarota, Chiara Soriani, Raphaelle Taub, Fiona Morgan, Jiro Sakai, Sarah L. Veatch, Clare E. Bryant, Pietro Cicuta

## Abstract

Signalling is of particular importance in immune cells, and upstream in the signalling pathway many membrane receptors are functional only as complexes, co-locating with particular lipid species. Work over the last 15 years has shown that plasma membrane lipid composition is close to a critical point of phase separation, with evidence that cells adapt their composition in ways that alter the proximity to this thermodynamical point. Macrophage cells are a key component of the innate immune system, responsive to infections, regulating the local state of inflammation. We investigate changes in the plasma membrane’s proximity to the critical point, as a response to stimulation by various pro- and anti-inflammatory agents. Pro-inflammatory (IFN-*γ*, Kdo-LipidA, LPS) perturbations induce an increase in the transition temperature of the GMPVs; anti-inflammatory IL4 has the opposite effect. These changes recapitulate complex plasma membrane composition changes, and are consistent with lipid criticality playing a master regulatory role: being closer to critical conditions increases membrane protein activity.

## Introduction

Macrophages are extremely versatile cells of the innate immune system able to activate and adapt their functionality depending on the specific milieu ***Martinez and Gordon (2014)***. Following phagocytosis of material resulting from trauma, or pathogens, or detection of specific functional molecules, macrophages can change their gene regulatory state and polarize into activated states, where for example they produce immune effector molecules such as cytokines for intercellular communication ***Taylor et al. (2005); Mosser and Edwards (2008); Brandsma et al. (2018)***. The responses manifested as a consequence of different stimulations have been classified in two broad activation states, based on both genetic expression profiling and phenotypic behavior: M1, or classically activated, macrophages have an enhanced bactericidal and tumoricidal capacity and produce high levels of pro-inflammatory cytokines, while M2 macrophages produce low levels of cytokines and have a wound-healing capacity by contributing to the production of collagen and extracellular matrix ***Martinez and Gordon (2014); Lawrence and Natoli (2011); Mosser and Edwards (2008)***. The stimuli that promote M1 macrophage activation are mainly IFN-*γ*, LPS and GM-CSF. IFN-*γ* is a cytokine mainly produced by natural killer (NK) and T helper 1 (Th1) cells; signaling from the IFN-*γ* receptor (IFNGR) controls the regulation of specific genes related to the production of cytokine receptors, cell activation markers and adhesion molecules ***Martinez and Gordon (2014)***. Lipopolysaccharides (LPS) are a class of molecules of the outer membrane of gram-negative bacteria, these molecules are recognized by the TLR4 receptor ***Park et al. (2009); Kawai and Akira (2010)***. TLR4 activation triggers the downstream production of pro-inflammatory cytokines such as TNF-*α* and presentation of antigens ***Akira and Takeda (2004)***. In contrast, macrophages polarize into M2 mainly in response to IL4 and IL13 stimuli. IL4 is produced by T helper 2 (Th2) cells, basophils, and mast cells in response to a tissue injury and in presence of some fungi and parasites ***Mosser and Edwards (2008)***. M2 cells are sensitive to infections, their production of pro-inflammatory cytokines is minimal, and their phagocytic activity is low ***Martinez and Gordon (2014); Mosser and Edwards (2008)***.

In the transduction of signals a fundamental regulatory role is thought to be played by the plasma membrane composition ***Simons and Toomre (2000)***. There are many examples of specific protein-lipid affinity, but also strong evidence of more general mechanisms such as the propensity of lipid mixtures to form cholesterol rich domains, or domains of a preferred thickness, which then imply a preferred partitioning of certain transmembrane proteins ***Simons and Sampaio (2011); Stone et al. (2017); Veatch and Cicuta (2018)***. Any mechanism that modifies local recruitment of membrane proteins, in the context of an assembly step such as dimerization necessary for function, can therefore directly be a regulator of receptor activity. This generalises a well known theme in membrane biochemistry, that proteins with lipid raft affinity have a higher chance to interact ***Pralle et al. (2000)***. The key structures in this study of macrophages, the TLR4 receptor and its co-receptor CD14, are both known to have raft affinity: CD14 is found in lipid rafts both before and after LPS activation, while TLR4 receptors are initially found in non-raft regions and then translocate to rafts after the activation ***Triantafilou et al. (2002)***. It has also been shown that the use of lipid raft inhibitors reduces significantly the production of cytokines related to LPS activation ***Nakahira et al. (2006)***. Moreover, lauric fatty acid seems to be responsible for the recruitment and dimerization of TLR4 into lipid rafts ***Wong et al. (2009)***. All together these facts strongly hint that plasma membrane composition, and in particular the propensity to form lipid rafts or domains, are fundamental regulators of protein interaction; we explore this theme with respect to activation of macrophages and the activity of TLR4 receptors.

Various authors have put forward the idea that the lipid raft phenomenology is linked to the propensity for the lipidic component of the membrane to undergo liquid-liquid separation ***Veatch and Cicuta (2018)***, as was observed in plasma membrane extracts ***Veatch et al. (2008)***. Vesicles extracted from the plasma membrane of cells have the same characteristics of certain ternary lipid mixtures, of particular interest the spontaneous appearance of transient lipid domains which is a universal property of systems in vicinity of a critical point ***Veatch et al. (2008); Honerkamp-Smith et al. (2008)***. From a biological point of view, being poised close to a critical point could be advantageous to accelerate a whole set of membrane biochemistry, since the cell would require much less energy to create lipid heterogeneity. Modulating the lipid composition is thus a mechanism for global regulation of activity on the membrane ***Veatch and Cicuta (2018)***. Giant plasma membrane vesicles (GPMVs) allow to study the properties of the membrane lipids as isolated systems ***Scott (1976); Scott and Maercklein (1979)***. These vesicles are thought to maintain the protein and lipid diversity of the mother membrane ***Scott and Maercklein (1979); Fridriksson et al. (1999)***, and at low temperatures the lipids can phase separate laterally into micron sized domains ***Baumgart et al. (2007); Veatch et al. (2008); Kaiser et al. (2009)***. GPMVs as systems to study the criticality of the plasma membrane have shown systematic dependency on growth temperature ***Gray et al. (2015)*** and cell cycle ***Burns et al. (2017)***, and on the epithelial-mesenchymal transition in cancer cells ***Tisza et al. (2016)***, and indeed in both situations the transition temperature of GPMVs recapitulates broad systematic composition changes that move the cell composition closer or farther from the critical point. In literature there are previous studies on the effect on lipid composition of macrophage activation ***Dennis et al. (2010); Andreyev et al. (2010)***, but these are bulk assays and report on the changes in a huge number of lipid species, making it difficult to interpret the results in simple terms. The work presented here shows that these complex changes in lipidomics may have a simple interpretation, in terms of their effect on the membrane phase separation. Investigating the effects of different kinds of macrophage cell stimulants (LPS, KLA, IFN-*γ*, IL4), known to differentiate macrophages into two different activation states, we show opposite changes with respect to the proximity of the critical point in the two cell types, consistent with biological function.

## Materials and methods

### Cell Culture

The immortalized BMDM cell lines were obtained from Dr. Eicke Latz (Institute of Innate Immunity at the University of Bonn, Bonn, Germany), and Dr. Kate Fitzgerald and Dr. Douglas T. Golenbock (University of Massachusetts Medical School, MA, USA). C57BL6 TLR4-/- mice were obtained from Dr. S.Akira (Osaka University, Osaka, Japan) ***Hoshino et al. (1999)***. iBMDM and TLR4^−/−^ iBMDM were maintained in Dulbecco’s Modified Eagle’s Medium (DMEM; Sigma-Aldrich, MO, USA) supplemented with 10% (v/v) heat-inactivated Hyclone fetal calf serum (FCS; Thermo Scientific, UT, USA), 2mM L-glutamine (Sigma-Aldrich), 100 U/mL penicillin and streptomycin (Sigma-Aldrich), and 20mM HEPES (Sigma-Aldrich). Cells are cultured for at least two days and brought to confluence in a single 175 cm^2^ flask. From confluence, cells are plated in separate dishes. To test the effect of stimulants on the melting temperature an equal number of cells are plated for each condition; we use a density of about 6-7·10^3^ cells/mm^2^. After 12 hours the culture medium is changed with (or without for the control condition) the addition of stimulating agents. Then, after the stimulation time, 12 hours, we start the GPMVs production protocol. Cell stimulating agents are used at the following concentrations: IFN-*γ* 20 ng/ml (PeproTech); LPS from *Salmonella* Typhimurium 10 ng/ml (Enzo Life Sciences); Kdo-LipidA 100 ng/ml (KLA, Avanti Polar Lipids); IL-4 20 ng/ml (PeproTech), and left for 12 or 24 hours. These doses where chosen according to previous work on M1/M2 macrophage differentiation ***Vats et al. (2006); Tatano et al. (2014) Kigerl et al. (2009)***.

To measure the T_*m*_ vs cell density dependency, density was measured in two different ways. For some experiments, images of the culture were acquired with a low magnification objective and the density estimated by counting cells from the image and then dividing their number by the field of view area. The same dish was then used to produce GPMVs immediately after. Otherwise for each density we had twin dishes, one was used to count the cells with the hemocytometer, whilst the other was used to produce GPMVs. To check the effect of stimulation on growth rate, an equal number of cells were plated in a multi-well then, for each condition (control, IL4, LPS); cells where counted with the hemocytometer after cell adhesion (0h), then stimulated, according to previously specified concentrations, and counted after 12h. Cells where initially plated to have about 6-7·10^3^ cells/mm^2^ at 0h.

### GPMVs production

The procedure for membrane labeling and GPMVs production follows the protocols in ***Sezgin et al. (2012)*** and ***Gray et al. (2013)***. The cells are gently washed twice with PBS, then DiI-C12(3) (Life Technologies) dye solution 50 *μ*g/ml in PBS is added and left on ice for 10 minutes to allow incorporation into the membrane. Then the cells are washed five times with PBS and twice with GPMV buffer. GPMV buffer is formed by 10mM HEPES, 150mM NaCl, 2mMCaCl_2_, the pH is adjusted to 7.4 with HCl or NaOH. Lastly the vesiculating agent is added and the cells are left in the incubator for 1.5 hours at 37°C. 20 *μ*l of vesiculating agent (2mM DTT, 25mM PFA) is used for each ml of GPMV buffer. The medium is gently harvested and transferred into a falcon tube. The sample is left at 37°C enough to let the blebs deposit on the bottom of the tube: for a volume of 4 ml, 24 hours are enough for the whole sample to sediment.

### Isolation of Lipids, and Gel-assisted vesicles formation

For the lipids isolation procedure we followed the Bligh and Dyer method ***Bligh and Dyer (1959)***: 1 ml of GPMVs sample is collected and moved to a vial. Then are added 3.75 ml of 1:2 chloroform and methanol mixture, 1.25 ml of chloroform and 1.25 ml of distilled water. After each step the solution is vortexed for 1 minute. At this stage the GPMVs burst and the components dissolve in the solution. The mixture is then centrifuged at 1000 RPM for 5 minutes. This makes the chloroform/methanol fraction deposit at the bottom of the tube, together with the lipids, while the aqueous and water soluble component is isolated at the top. Proteins are preferentially located at the interface between the two phases. The bottom phase is then collected and left under vacuum to let the solvents evaporate. Finally lipids are redissolved in 100 *μ*l of chloroform.

The vesicles are produced through the gel-assisted method as described in ***Weinberger et al. (2013)***: 200 *μ*l of 5% (weight/weight) PVA solution is spread on a microscope coverslip with the help of a spincoater and then left to dry in an oven at 50°C for 30 minutes. Lipids dissolved in 100 *μ*l of chloroform are then spread on PVA gel. A chamber is formed with the help of a spacer and a second coverslip and filled with a solution of sucrose. After 30 minutes the vesicles are collected and diluted in glucose solution to allow vesicle sedimentation.

### Imaging

The samples are imaged on a Nikon Eclipse Ti-E inverted epifluorescence microscope using a Nikon PLAN APO 40× 0.95 N.A. dry objective and a IIDC Point Grey Research Grasshopper-3 camera. The perfect focus system (Nikon) maintains the sample in focus even during thermal shifts. The temperature of the sample is controlled with a home-made computer-controlled Peltier device. A thermocouple is placed in direct contact with the sample chamber. In each position a z-stack of 8 images is acquired, spanning across a range similar to the bleb size. The temperature is decreased across the whole sample with a ramp from 37 to 3°C in steps of 2°C; at each step the temperature is let to equilibrate for 15 seconds. The abundance of GPMVs produced can vary from cells prepared in different days, but usually from a dish of 5.5 cm diameter with confluent cells it is possible to produce blebs for at least 2 experiments. With the quantities described above, we are able to image up to 100-200 blebs in each field of view.

### Software processing

A custom Matlab software pipeline has been developed to automatically detect the position and radius of the GPMVs in the images. It uses the Hough transform to detect circular features. Then with the help of a graphical user interface the blebs are shown to the user one at the time, the user can interactively scroll the z-stack and decide if the bleb shows (a) a single phase, (b) phase coexistence or (c) unclear phenotype. The software randomly picks the vesicle to show, from the database of all the temperatures, i.e. in this stage the information about the temperature is kept hidden to the user, so that the decision process (assigning the type a/b/c) is unbiased.

## Results

Following established protocols, GPMVs are produced from macrophage cells using PFA and DTT. The sample is observed under an optical microscope and it is in contact with a temperature control system. The temperature is lowered from 37 to 3°C in steps of 2°C. At high temperatures all the vesicles show a uniform phase. Around 12-22°C phase separation domains start to appear in some GPMVs, and at low temperatures most of the GPMVs are phase separated (see Figure 1 A). For each temperature we calculate the fraction *f* (*T)* of GPMVs which show uniform phase or phase separation. Before producing GPMVs, macrophages are stimulated with one of IFN, LPS or KLA for 12 hours to induce pro-inflammatory response. In each data set (Figure 1 B-E) we compare the stimulated condition with its unstimulated control data set, since we noticed (as has been already reported in different cell types ***Gray et al. (2013***, 2015)) a significant variability in the transition temperature of independent repeats; in contrast, the transition temperatures of GPMVs from the same cultures, even split into separate dishes, are tightly distributed.

**Figure 1.**
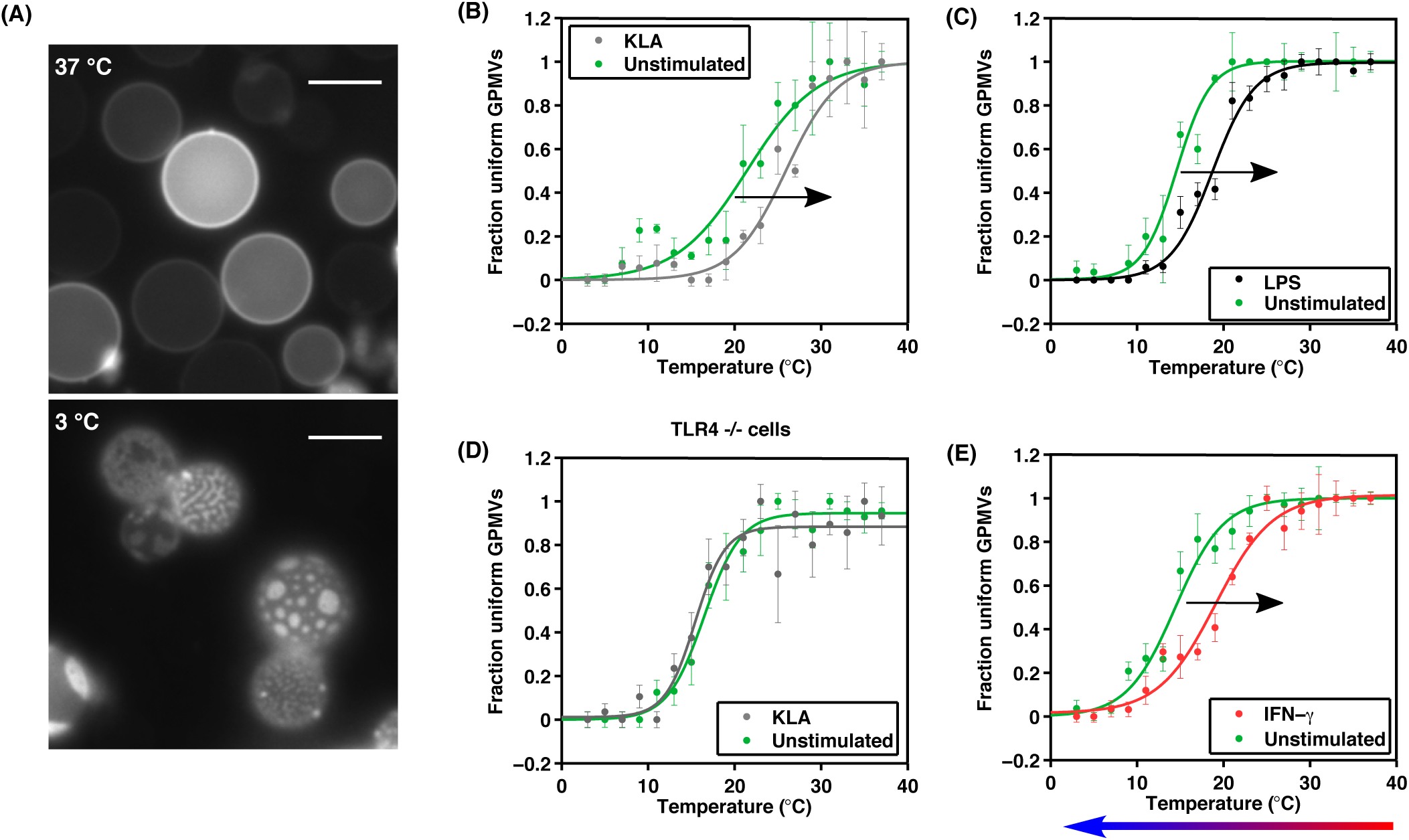
The plasma membrane of macrophage cells is poised close to phase separation, and the proximity to the critical point varies in response to signaling molecules. (A): Fluorescence microscope image of GPMVs at 37 and 3°C. Scalebar 5*μ*m. (B-E): Fraction of GPMVs showing just one phase over the total of vesicles observed in function of the temperature. The data show a sigmoidal trend and are fitted with a hyperbolic tangent from which are extracted the transition temperature at mid height and the width of the transition. We compare the sample obtained from cells treated with KdoLipidA (B), LPS (C), IFN-*γ* (E), for 12h compared to a non-treated control condition prepared in parallel. All these “pro-inflammatory” treatments shift the transition temperature towards higher temperatures. The colored arrow at the bottom indicates the direction of the temperature variation imposed on the GPMV samples during the imaging process. (D): The knock-out TLR^−/−^ cells do not vary transition temperature when stimulated with KdoLipidA (in contrast to panel (A)), remaining the same to the unstimulated controls.

The transition temperature *T*_*m*_ is obtained by fitting the *f* (*T)* data with an empirical sigmoidal curve:

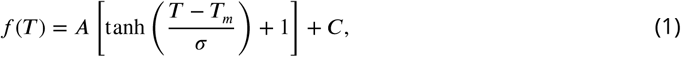

where *T*_*m*_ and *σ* are the most interesting parameters to describe the mean and the cell-to-cell variability (GPMVs originate from individual cells) in the transition temperature of the population. Error bars are associated with data points by randomly separating the measurements for a given temperature into three groups, and treating these as independent data sets.

Figure 1 B shows the effect of the cell stimulation with KLA for 12 hours. The comparison with the control condition shows a shift of 4.5°C in the GPMVs transition temperature to higher temperatures. As expected, LPS and KLA stimulations produce similar effects (see Figure 1 B-C). Indeed Kdo-LipidA is the active sub-unit of the LPS molecule which is recognized by the TLR4 trans-membrane receptor ***Park et al. (2009)***. Notice that the comparison between LPS and KLA has to remain qualitative since there isn’t a first-principles way to correlate the doses, except for the effects on activating cells. Both doses employed here are known to be able to saturate the cell response, for example in terms of TNF*α* production ***Andreyev et al. (2010); Vasan et al. (2007)***. As a control, repeating the same experiment of KLA stimulation, this time on TLR4^−/−^ macrophage cells, we obtained compatible transition trends (Figure 1 D) between the stimulated and unstimulated condition. This confirms that the observed temperature shift is originated from the metabolic change as a downstream effect of triggering NF-*κ*B signalling, more than any other spurious effect. We then stimulated cells with IFN-*γ*, known, like LPS, to have pro-inflammatory effects ***Martinez and Gordon (2014)***, and obtained the same qualitative effect on the plasma membrane transition temperature (Figure 1 E).

Since all the experiments with the “classically activated” conditions were showing a consistent shift in the same direction, we decided to stimulate the cells with interleukin-4 (IL4) which is known to induce a different type of differentiation ***Mosser and Edwards (2008)***. Macrophages treated with IL4 have different phenotypes and markers compared to the M1, and a different role in the immune response: they don’t produce pro-inflammatory cytokines, but suppress destructive immunity, and are involved for example in wound healing response ***Mosser and Edwards (2008)***. The curves in Figure 2 correspond to the control condition and to 24 hours of IL-4 stimulation. Also in this case the stimulation produces a temperature shift, but in contrast to the “classically activated” cells, in this case the *T*_*m*_ shifts towards lower temperatures.

**Figure 2.**
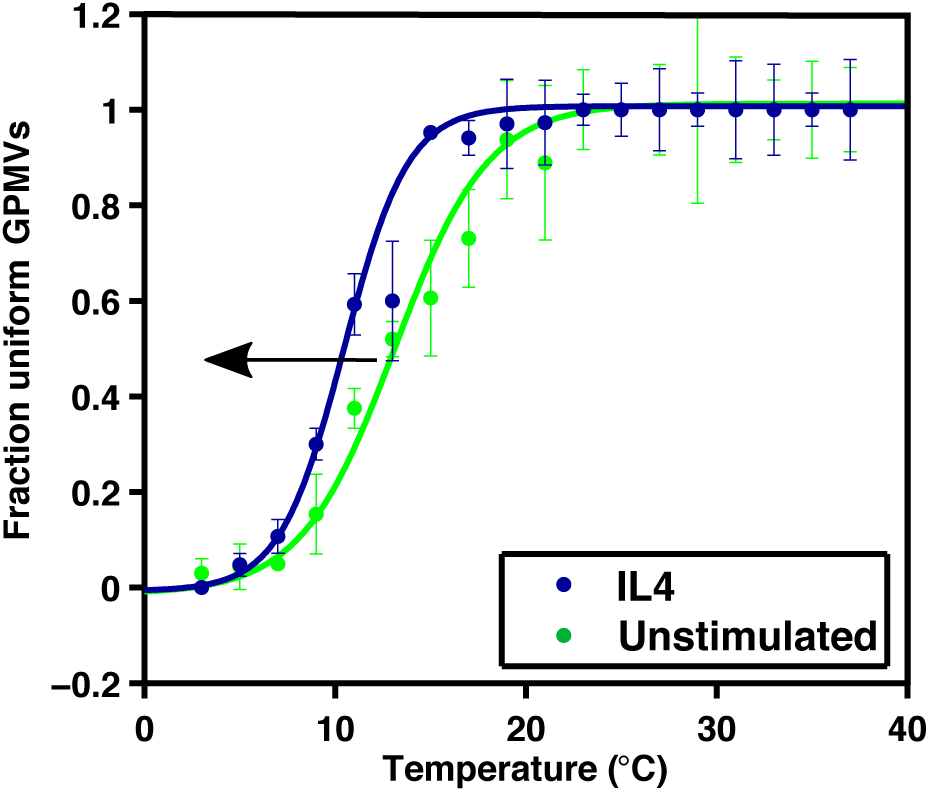
Anti-inflammatory treatment moves the composition of the plasma membrane farther from the critical point. The data show the fraction of uniform GPMVs as the temperature of the sample is varied. The two curves correspond to 24 hours of IL4 stimulation and to unstimulated conditions. T_*UNST*_ = (13.11 ± 0.49)°C, T_*IL*4_ = (10.46 ± 0.33)°C.

Collecting together all the *T*_*m*_ values from different stimulation experiments (see Figure 3 A) we can see how the IL4 data and the IFN-*γ*/LPS/KLA data are in two separate temperature ranges, with no data overlapping, while the values from the unstimulated experiments have a much wider range. Statistical analysis confirms the distributions to be significantly different (p<0.05) for almost all of the conditions. Calculating the temperature differences *T*_*stim*_ - *T*_*unstim*_ for all the 12 hours stimulation experiments (i.e. comparing with same day controls), the temperature shifts tighten (Figure 3 B) and show very consistent behaviors: the IL4 data points are all negative, whereas the others are all positive.

**Figure 3.**
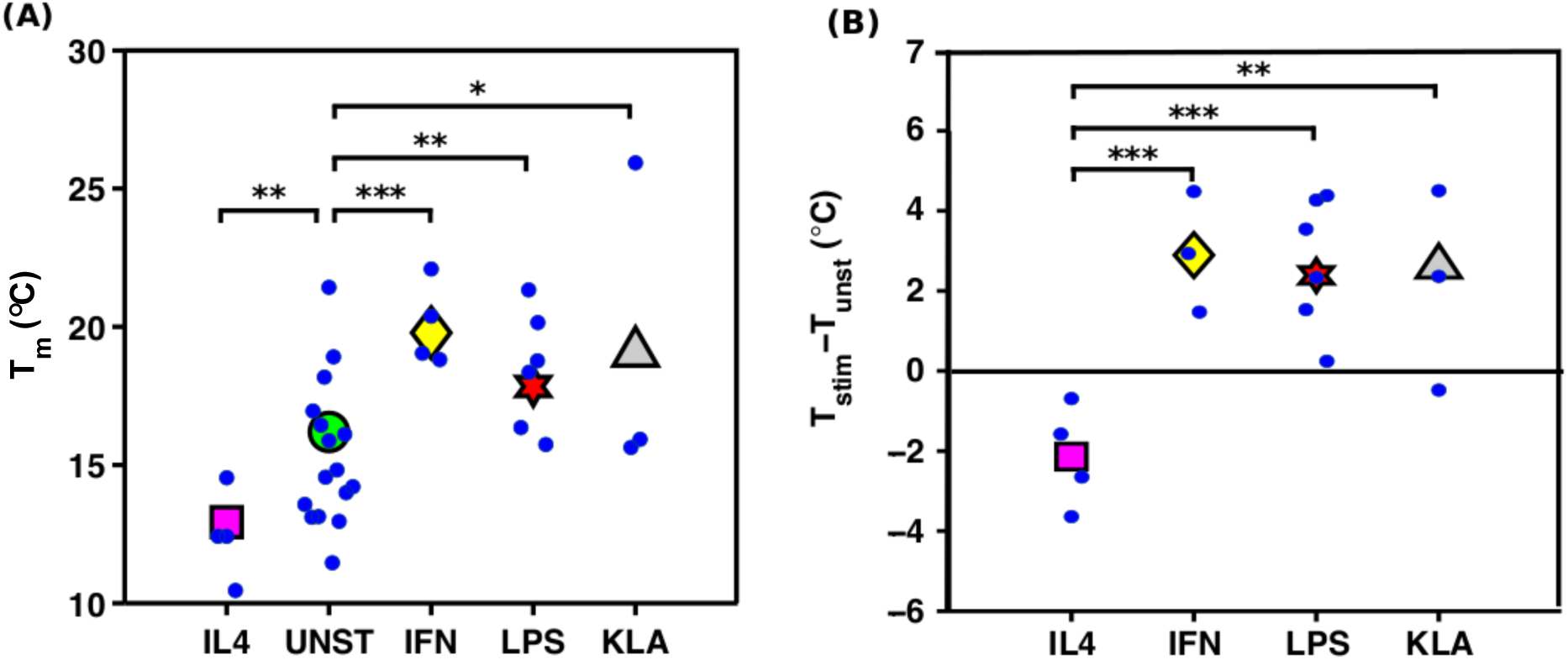
Pro- and anti-inflammatory treatments affect the transition temperature systematically. The scatter in the absolute transition temperature (particularly notable in the unstimulated UNST cells) is reduced significantly comparing with same-day unstimulated controls. (A): Fitted transition temperatures of vesicles produced by macrophage cells treated with IL4, IFN-*γ*, LPS or Kdo-LipidA for 12 hours. Each small data marker comes from an experiment with between 300 and 600 vesicles. The large markers indicate the average in each distribution, weighted with the errors on *T*_*m*_. (B): Temperature difference of each stimulation experiment with its control condition. From one-way analysis of variance (ANOVA) we obtained the distributions differences to be statistically significative with * p<0.1, ** p<0.05, *** p<0.005.

We then investigated cell density as one of the possible causes for the large variability of *T*_*m*_ in the control condition. For the experiments described above, we used a celluar concentration from about 6 to 7·10^3^ cells/mm^2^. The effectiveness of intracellular communication indeed depends on the cell density, and can be conveyed through both mechanical or chemical interaction ***Stow et al. (2009); Fortes et al. (2004); Lim et al. (2011)***. In Figure 4 we report the results of experiments done growing cells in a common flask, and then plated at different densities. The transition temperature correlates with the cell density, so that the curve that corresponds to the most crowded sample is on the right of the less dense samples (see Figure 4 A). Summarizing all the density measurements, we obtain a linear trend of the miscibility temperature as a function of the cell density (Figure 4 B). A similar trend has been obtained recently in similar experimental conditions for other cell types ***Gray et al. (2015)***, and possible causes are presented in the discussion.

**Figure 4.**
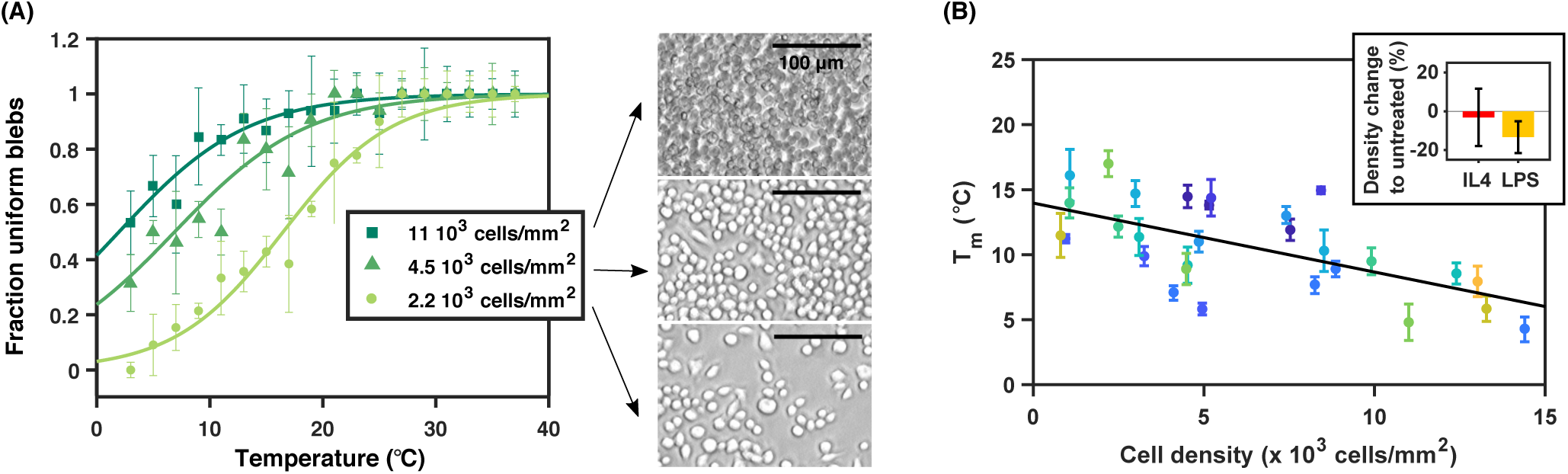
Cell crowding affects the phase transition of GPMVs. (A) There is a consistent shift in the data for the fraction of uniform GPMVs, from cell cultures at different densities. T_11_ = (4.8 ± 1.4)°C, T_4.5_ = (8.9 ± 1.2)°C, T_2.2_ = (17.0 ± 1.0)°C. (B) The miscibility temperatures obtained from the sigmoidal curve fitting, as a function of the cell crowding, showing that the denser samples have lower transition temperature. Same colors indicate repetition of the experiment on same day. The linear fit *y* = *a* + *cx* gives c = (−0.53 ± 0.26)°C mm^2^/cells and a = (14.0 ± 1.9)°C. In the sub-panel is represented the density change as effect of 12 hours of stimulation compared to an untreated sample at our typical experiment densities (7 · 10^3^ cells/mm^2^). Numerical values obtained over 4 repetitions are: (−3 ± 15)% for IL4 treated, and (−13 ± 8)% for LPS treated.

We had at this point to check the possibility that the temperature shift observed as a function of the pro/anti-activation treatment might be an indirect effect, due to a differential stimulus-dependent growth. To check for this, we measured cell growth through the difference in cell density after IL4 and LPS stimulation. Reproducing our typical experimental conditions (7·10^3^ cells/mm^2^, and 12 h stimulation), we obtained a non-significant change in the density of IL4 treated cells compared to the unstimulated condition, while the LPS showed a decrease of 13%. Putting together the growth rate reduction with LPS with the calibrated cell-concentration results, for the LPS condition we obtain (as an indirect effect of the stimulant on the cell culture growth rate) an expected change of the melting temperature of about -0.5°C compared to the untreated condition. Therefore this important control shows that the ∼ 2 degrees difference in T_*m*_ seen between untreated and LPS stimulated is due only in small part to cell density, so most of the effect has to be accounted for by processes independent of density, downstream of the LPS signalling pathway.

The high quality imaging allowed us to investigate also the shape of the phase separation domains appearing in blebs at low temperatures. Some of the domains indeed appear to have an irregular rigid shape similar to a gel phase domain, while others look more rounded like in the situation of liquid-liquid coexistence. With the help of a graphical user interface that shows a 3-4 frames time sequence of the vesicle, GPMVs with irregular domains where identified as the one presenting rigid and not rounded dark regions (see Figure 5-supplement 1). The appearance of gel looking domains on GPMVs has already been reported ***Gray et al. (2015)***, but this is the first attempt for a quantification of the phenomenon. Three sets of data in different conditions are shown in Figure 5. In all the cases, in spite of the noise, the fraction of irregular domains over the total of phase separated GPMVs has a clear growth at low temperatures, reaching about 0.4 at 3°C. On the other hand we don’t see any significant difference in these trends comparing different stimulation conditions. In the event that these irregular domains could be confirmed as gel domains, this kind of analysis would provide an additional piece of information on the phase diagram of the biological membrane lipid mixture (on which we don’t have almost any knowledge) and might be particularly important in cell biology regulation involving cholesterol ***Ayuyan and Cohen (2018)***.

**Figure 5.**
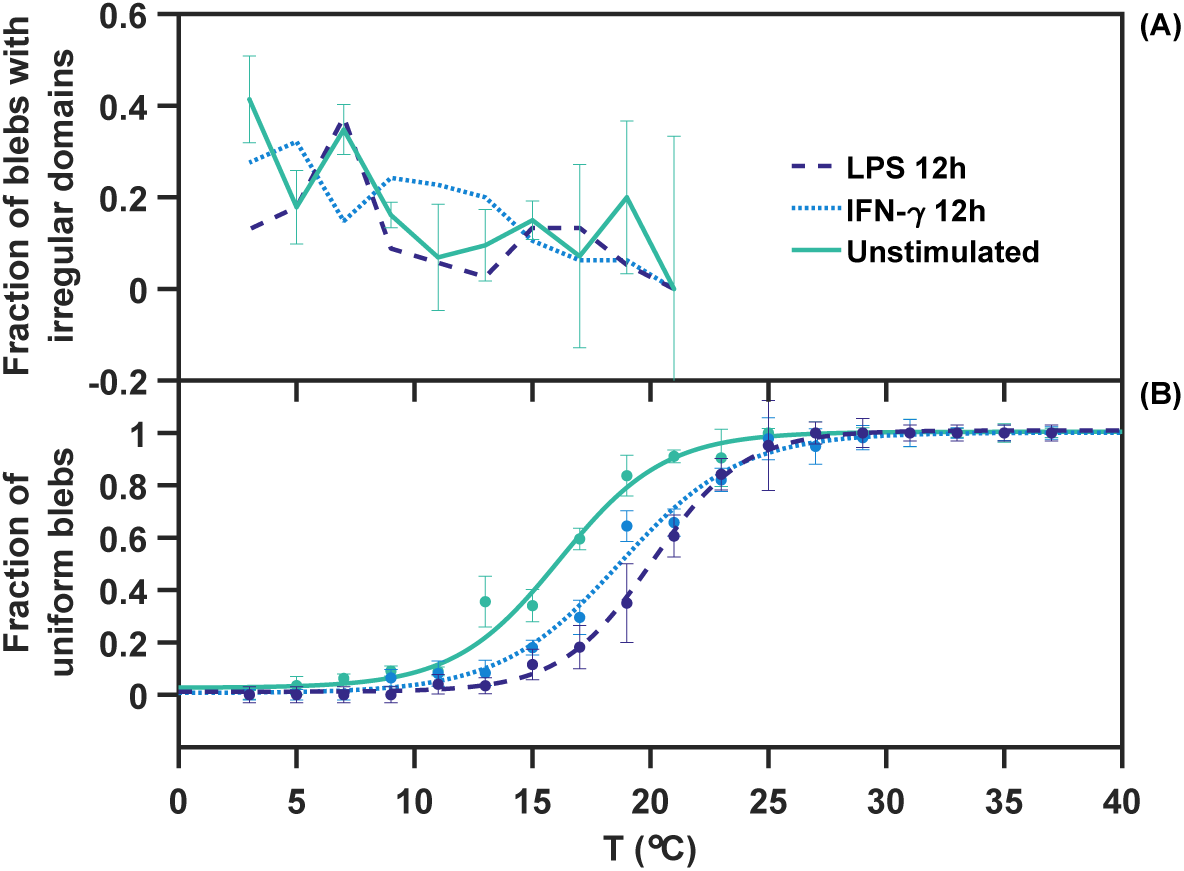
At very low temperatures, irregular shaped domains are observed and attributed to a gel phase. (A) The fraction of GPMVs with irregular domains (over the total of phase separated GPMVs) increases at low temperatures. This fraction grows below *T*_*m*_, as can be seen comparing in (B) the ‘conventional’ data on liquid-liquid phase separation for the three conditions indicated in the legend.

The experiments described so far provide evidence that the composition of the plasma membrane is regulated according to the external milieu, but we still don’t know if this change involves just the lipids and/or also the membrane protein composition or abundance. To address this, we performed an important and seldom considered control: comparing the melting temperature of GPMVs with the same sample after a lipid purification process. The GPMVs sample was divided in two aliquots, and one of them was dissolved, purified and the vesicles re-formed trough the gel-assisted formation technique, as described in the methods section. The two samples are found to have compatible values for their miscibility temperature (see Figure 6 and Figure 6-supplement 1), meaning that the phase separation phenomenon on GPMVs is lipid driven and that the miscibility temperature is mostly unperturbed by membrane proteins. It is also worth remarking that these reconstituted vesicles have lost any possible bilayer asymmetry maintained in the GPMVs.

**Figure 6.**
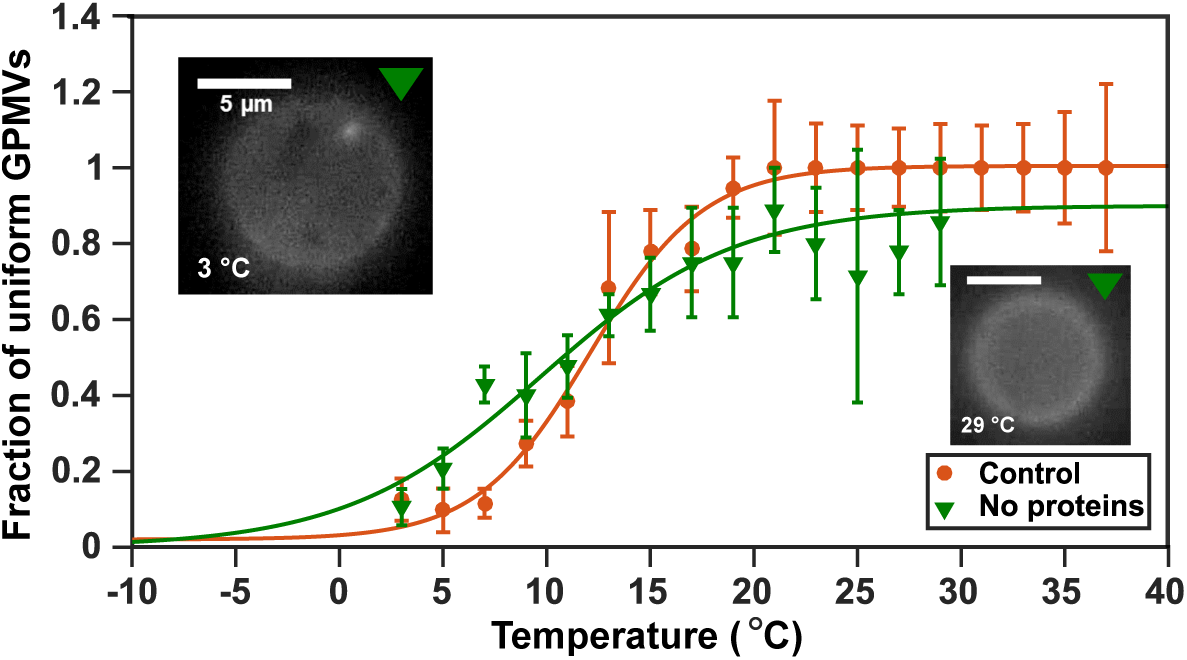
A key control with purified lipids excludes the role of proteins in the phenomena reported above. Comparison between the distribution of uniform GPMVs against analysis of vesicles re-formed from the just the lipidic component purified from the same sample. The melting temperature is compatible, within the error, meaning that the lipids are unperturbed in the determination of the melting temperature. T_*noproteins*_ = (11.1±1.4)°C, T_*control*_ = (12.2±0.5)°C.

## Discussion

It is well known that the plasma membrane is not just a passive support for activity by membrane proteins, and here we have developed the theme that the property of lipids to phase segregate relates to protein interactions ***Veatch and Cicuta (2018); Kimchi et al. (2018)***. GPMVs are an extremely useful system to understand this aspect of plasma membranes because they maintain the composition of the original membrane, but they can be studied as an isolated structure and subjected to stringent controls. Our results add to the body of evidence that proximity to the critical point for phase separation can be a global regulator favoring activity, and for the specific case of macrophage cell activation our results are consistent with previous findings that the early stages of TLR4 activation take place in raft domains ***Triantafilou et al. (2002)***.

### Effect of stimulation on plasma membrane transition temperature

We have seen how treatment of macrophages with different stimulating agents affects the melting temperature of GPMVs. All the stimulants used (IFN-*γ*, LPS, KLA, and IL4) induced a shift of few degrees compared to the control condition, meaning that in all the cases the membrane composition has changed as a consequence of the activation of specific signaling pathways. Moreover, IFN-*γ*, LPS and KLA increased the transition temperature (*T*_*m*_), whereas IL4 had the opposite effect decreasing *T*_*m*_. Given that the first three stimulants can be connected to the activation into the M1 state in macrophages, whereas IL4 is responsible for the differentiation into the M2 state, this result sheds new light on the importance of plasma membrane composition in the immune response, and suggests new ways in which lipidomics may be involved in the regulation of the host defense strategy.

From the point of view of the membrane composition, if the melting temperature increases (coming closer to physiological temperature) it means that spontaneous lipid domains are longer lived and larger, so that membrane components can partition more strongly; also, the energy cost to recruit a particular lipid micro-environment around a protein is reduced ***Kimchi et al. (2018)***. It has been calculated that due to this universal phenomenon, the proximity to critical point, spontaneous lipid domains exist at sizes of around 22 nm for GPMVs from RBL cells ***Veatch et al. (2008)***. This argument considers the dimension of the correlation length *ξ* at a physiological temperature (T = 37°C), and experiments that measured *T*_*m*_, then using the expression *ξ* = *ξ*_*o*_*T*_*m*_/(*T* − *T*_*m*_) ***Honerkamp-Smith et al. (2008)***. This same argument can now be extended, in light of the results presented here: keeping the same value of *ξ*_*o*_ from ***Veatch et al. (2008)*** (because this is a quantity liked to the size of the lipid) we can estimate the effect of an increase in *T*_*m*_ from *T*_*m M*1_ = 13 to *T*_*m M*2_ = 20 °C (as from figure 3). This results in an increase of the correlation length of the order of 40% (from *ξ*_*M*1_ = 12 to *ξ*_*M*2_ = 17 nm). We expect this to have effect on confinement of proteins and their local concentration, and thus to affect for in the particular system here the balance of dimerisation in TLR4 receptors, and hence regulate the initiation of signaling pathways.

### Speculative correlation between membrane composition and receptor activity

We suggest here a possible correlation between the role of the cell in the immune defense and the changes in its membrane composition. One can imagine that these cells, depending on their activation state, regulate their lipid composition in such a way to tune the proximity to the critical point, and hence in turn the typical dimension and lifetime of spontaneous lipid domains, in order to be more or less reactive towards external stimuli. An M1 cell would have bigger and more long-lasting lipid domains, leading to increased activity of TLR4 receptors, which have raft affinity ***Plóciennikowska et al. (2014); Triantafilou et al. (2002); Pfeiffer et al. (2001); Triantafilou et al. (2004)*** (e.g. by increased recruitment to the membrane, and increased dimerization), to induce a faster and stronger inflammatory response with consequent production of inflammatory cytokines. In contrast, in M2 cells the activation of the TLR4 to NF-*κ*B pathway would be down-regulated through the lipid composition effect. An important element in support of this hypothesis is the reported increased sensitivity to LPS after IFN-*γ* treatment, both in mice ***Matsumura and Nakano (1988)*** and in macrophages in vitro ***Darmani et al. (1994)***, where a 66% increase of the LPS binding efficiency has been measured.

### Effect of cell density on *T*_*m*_

We see a shift of *T*_*m*_ with cell density. One could relate this result with the shift given by the different kind of stimulations, venturing a picture in which the overcrowded populations have some common behavior with M2 cells. In this picture, the crowded populations, with no need to further recruit cells and promote additional inflammation against possible infections, diminish their cytokine production, thus acting more like M2 cells. This hypothesis is supported by the observation that BMDMs from high density cultures secrete less pro-inflammatory cytokines and have lower phagocytic ability ***Lee and Hu (2013)***. Moreover the number of cells showing typical M2 membrane markers like CD11c and Ly-6CLy-6G increases in dense cultures ***Lee and Hu (2013)***.

A second mechanism could be identified in the asymmetry in the exposed membrane at different densities. Indeed at high densities the only region of the cell exposed is the top surface while the neighbor cell hide the lateral sides. The vesicles forming from the free surface could therefore be affected by the differences in the membrane composition due to the basal/apical polarization. To test the hypothesis of the interaction through cytokines, we performed an experiment where the medium was periodically changed every 2 hours. This treatment did not produce a significative change in *T*_*m*_ (see Figure in Supplementary Materials), meaning that the density effect is mainly due to interaction through mechanical contact.

Even though the density has been proven to be an important factor in the day-to-day variability in the *T*_*m*_ of unstimulated macrophages, this is not enough to explain the variability between independent repeats, indeed just keeping the cells in separate cultures is enough to produce some variability (Figure 4-supplement 1). To investigate the cause of the *T*_*m*_ day-to-day variation, the effect of cell density was tested, and the results showed that denser populations gave a lower *T*_*m*_ in GPMVs. The same experiment has been very recently performed on rat basophilic leukemia cells (RBL) ***Gray et al. (2015)*** with the same outcome, the authors suggesting that dense populations could have different physical membrane properties to be able to sense and communicate with touching cells ***Frechin et al. (2015)***. Our hypothesis is that cell density indirectly induces a decrease in T_*m*_, perhaps by triggering the production of cytokines with the same effect of IL4. This picture is supported by a study in which M1/M2-like differentiation was induced by the population density ***Lee and Hu (2013)***. Indeed, among other things, denser cell cultures where found less efficient in the production of cytokines than sparser ones after LPS stimulation ***Lee and Hu (2013)***. We tested this hypothesis comparing the T_*m*_ of two samples plated at the same density. GPMVs where produced after 12 hours, but during this time in one of the samples we changed the medium every 2 hours. This “washed” sample shows a higher T_*m*_ compared to the control, where cytokines would be accumulating in the medium, see Figure 4-supplement 2. This is compatible with a scenario in which the control condition is affected by an accumulation of M2-inducing cytokines such as IL4.

## Conclusions

For the first time, vesicles from purely the lipid fraction of GPMV from plasma membrane of macrophages have been observed, and their phase behaviour compared to the GPMV. This showed that both form phase separated domains on cooling, that the composition is close to a critical point, and that the melting temperature is unaffected by the presence of proteins. Also for the first time, we quantified the fraction of irregular domains on GPMVs, observing an increase of these at low temperatures. These are likely to be general results common to plasma membrane compositions in many cell types. The main biological question addressed here concerns macrophage cells, which we conditioned via pro- and anti-inflammatory stimuli, before extracting GPMV and measuring the phase transition temperatures. Considering all the transition temperatures together we get a very consistent picture: transition temperatures following IL4, as opposed to IFN-*γ*/LPS/KLA treatment, form two non-overlapping intervals (respectively at 10-15°C and 15-25°C). The absolute temperature changes induced by stimulation are always around 2°C compared to control. We have described a physical mechanism that can underpin this correlation between the immune response role of macrophage cells and the lipid composition of their plasma membranes, where signaling activation initiates. Much remains to be discovered within the ‘critical lipidomics’ paradigm, specifically direct experiments are becoming possible thanks to superresolution approaches ***Stone et al. (2017); Veatch and Cicuta (2018); Brandsma et al. (2018)***, probing membrane protein copy numbers and states of aggregation and how these are affected by the proximity to lipid mixture critical points.

## Acknowledgments

Research was funded by EU Marie Curie action ITN TransPol (EC), NIH-R01GM110052 and NSF-MCB1552439 (SLV), Cambridge University Commonwealth, European and International Trust (JS) and ITN BioPol (PC), Wellcome Trust Investigator grant 08045/Z/15/Z (CEB).

---

**Table 1.**
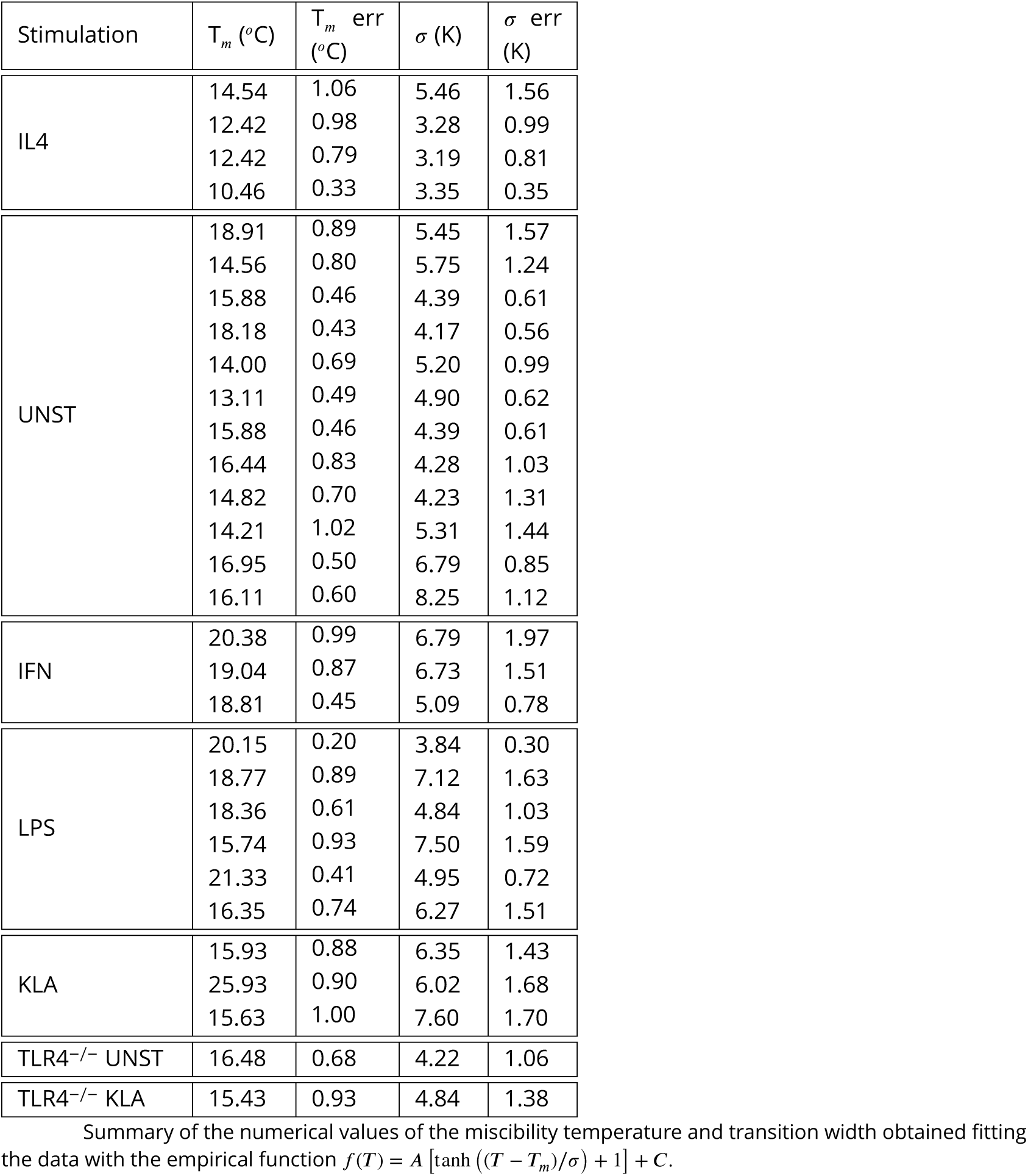
Summary of the numerical values of the miscibility temperature and transition width obtained fitting the data with the empirical function *f* (*T)*= *A* [tanh (*T* − *T*_*m*_)/*σ* + 1] + *C.*

| Stimulation | $T_m$ (°C) | $T_m$ err (°C) | $\sigma$ (K) | $\sigma$ err (K) |
| --- | --- | --- | --- | --- |
| IL4 | 14.54 | 1.06 | 5.46 | 1.56 |
|  | 12.42 | 0.98 | 3.28 | 0.99 |
|  | 12.42 | 0.79 | 3.19 | 0.81 |
|  | 10.46 | 0.33 | 3.35 | 0.35 |
| UNST | 18.91 | 0.89 | 5.45 | 1.57 |
|  | 14.56 | 0.80 | 5.75 | 1.24 |
|  | 15.88 | 0.46 | 4.39 | 0.61 |
|  | 18.18 | 0.43 | 4.17 | 0.56 |
|  | 14.00 | 0.69 | 5.20 | 0.99 |
|  | 13.11 | 0.49 | 4.90 | 0.62 |
|  | 15.88 | 0.46 | 4.39 | 0.61 |
|  | 16.44 | 0.83 | 4.28 | 1.03 |
|  | 14.82 | 0.70 | 4.23 | 1.31 |
|  | 14.21 | 1.02 | 5.31 | 1.44 |
|  | 16.95 | 0.50 | 6.79 | 0.85 |
|  | 16.11 | 0.60 | 8.25 | 1.12 |
| IFN | 20.38 | 0.99 | 6.79 | 1.97 |
|  | 19.04 | 0.87 | 6.73 | 1.51 |
|  | 18.81 | 0.45 | 5.09 | 0.78 |
| LPS | 20.15 | 0.20 | 3.84 | 0.30 |
|  | 18.77 | 0.89 | 7.12 | 1.63 |
|  | 18.36 | 0.61 | 4.84 | 1.03 |
|  | 15.74 | 0.93 | 7.50 | 1.59 |
|  | 21.33 | 0.41 | 4.95 | 0.72 |
|  | 16.35 | 0.74 | 6.27 | 1.51 |
| KLA | 15.93 | 0.88 | 6.35 | 1.43 |
|  | 25.93 | 0.90 | 6.02 | 1.68 |
|  | 15.63 | 1.00 | 7.60 | 1.70 |
| TLR4<sup>-/-</sup> UNST | 16.48 | 0.68 | 4.22 | 1.06 |
| TLR4<sup>-/-</sup> KLA | 15.43 | 0.93 | 4.84 | 1.38 |

**Figure 4–Figure supplement 1.**
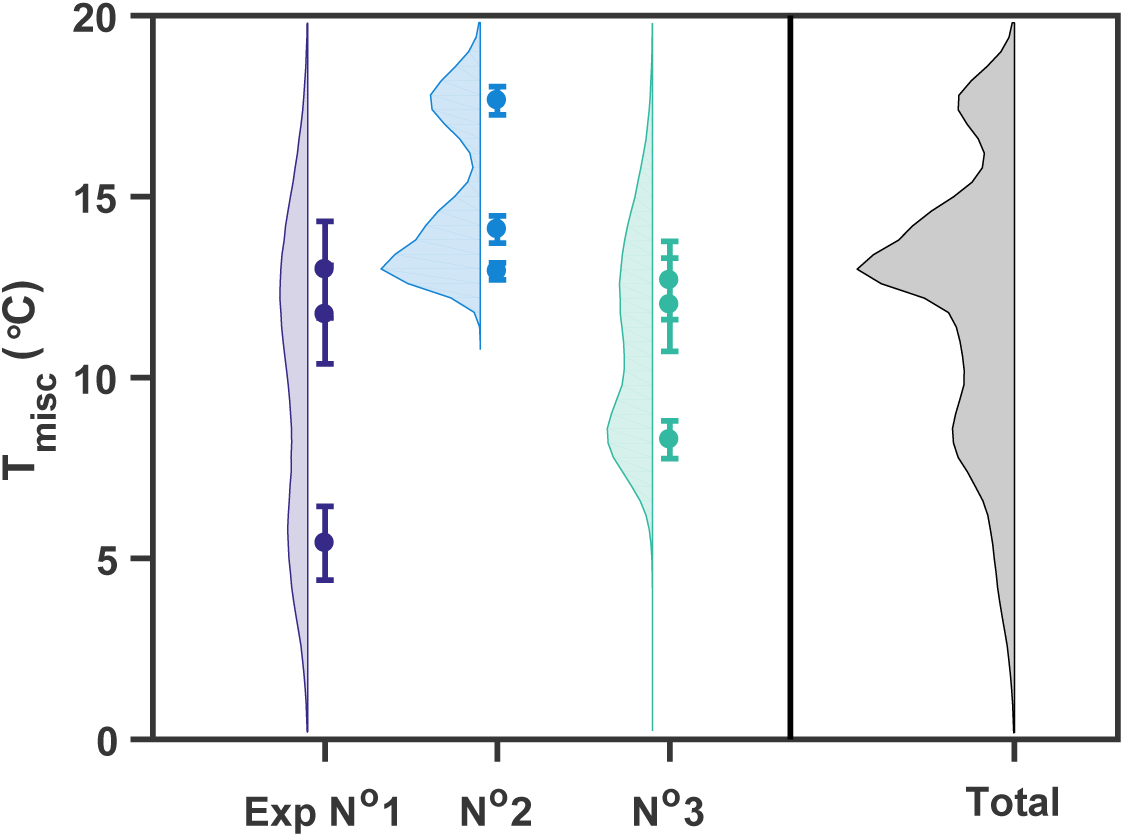
Miscibility temperature of three repetitions of the same experiment in which macrophage cells at the same density are cultured in three separate flasks as replicates for two days. The data points have a wide temperature range both within the replicates and comparing the three experiments. Data points are also interpreted as representative of a whole distribution with the same average and standard deviation. In spite of the large variation, the sum of the distributions shows a main peak around 13°C. Continuous distributions are obtained simulating gaussian distributed numbers with mean and standard deviation equal to the value of the data and their error.

**Figure 4–Figure supplement 2.**
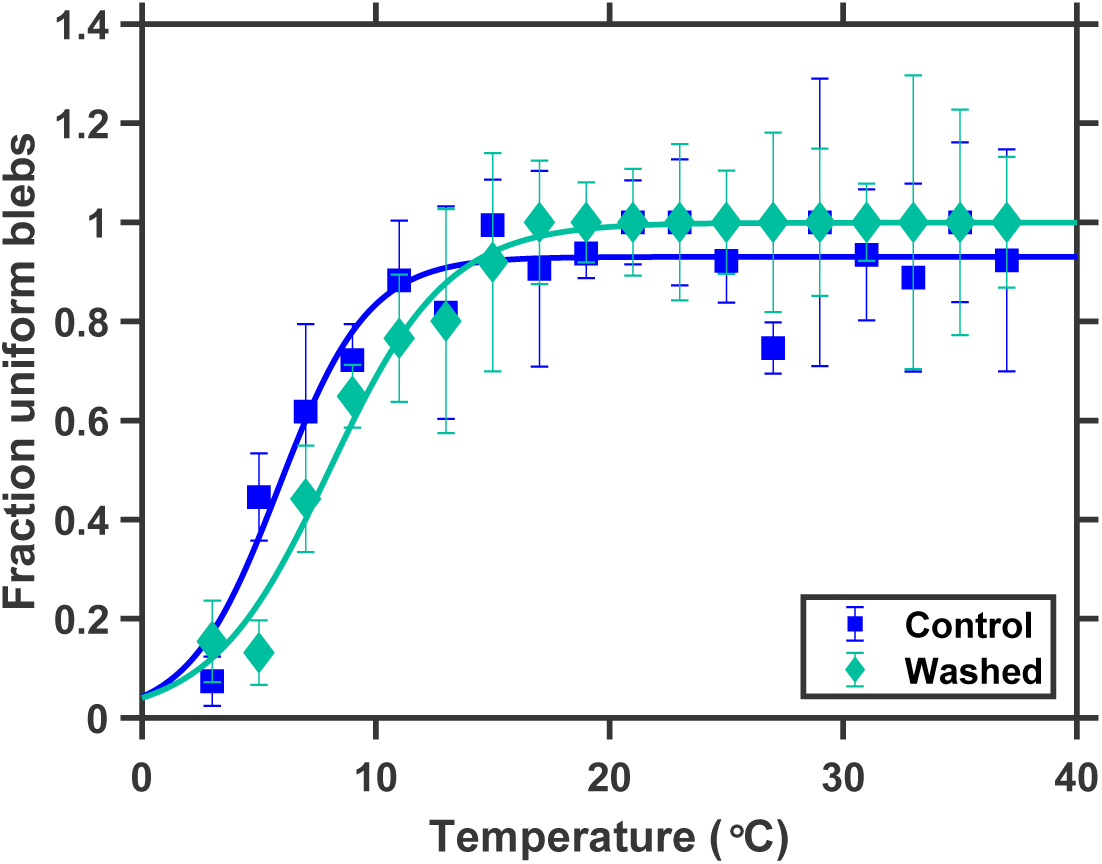
Controll experiment to test the effect of intracellular comunication through secretion of cytokines on the melting temperature of GPMVS. The two samples where plated at the same density, and blebbing was induced after 12 hours during which the medium of the “washed” sample was changed every two hours. The control sample shows a lower T_*m*_ compared to the “washed”, see Figure 4-supplement 2. This is compatible with a scenario in which the control condition is affected by un accumulation of M2-inducing cytokines like IL4. T_*contol*_ = 5.8 ± 1.0, T_*washed*_ = 7.9 ± 1.2

**Figure 5–Figure supplement 1.**
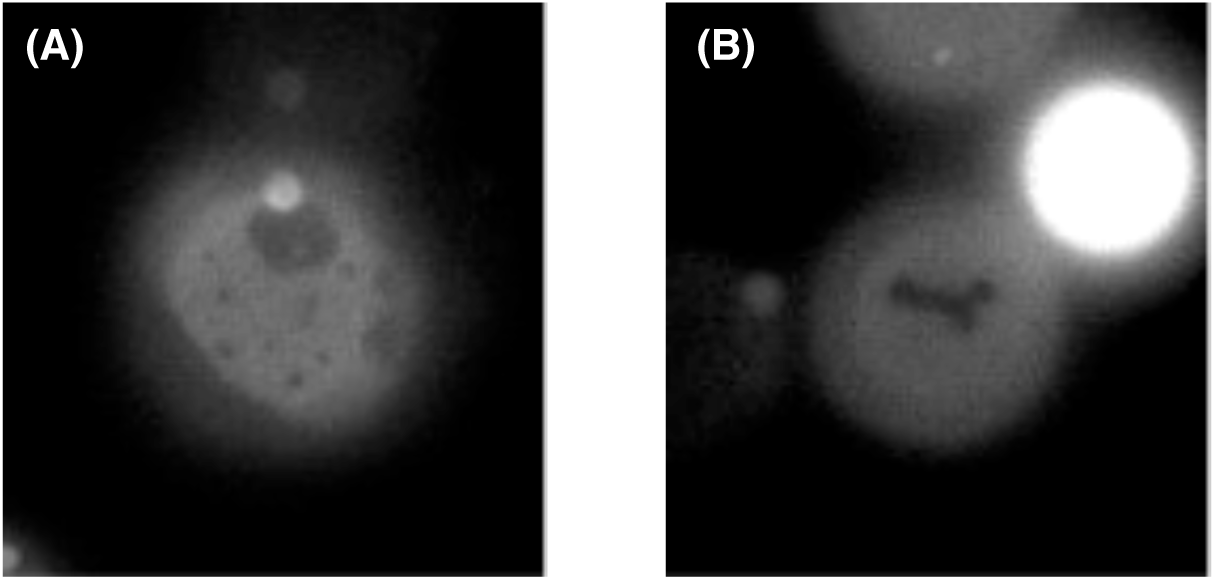
Example GPMVs showing round domains (A) and an irregular domain (B). These different shapes likely correspond respectively to liquid-liquid and liquid-gel phase coexistence.

**Figure 6–Figure supplement 1.**
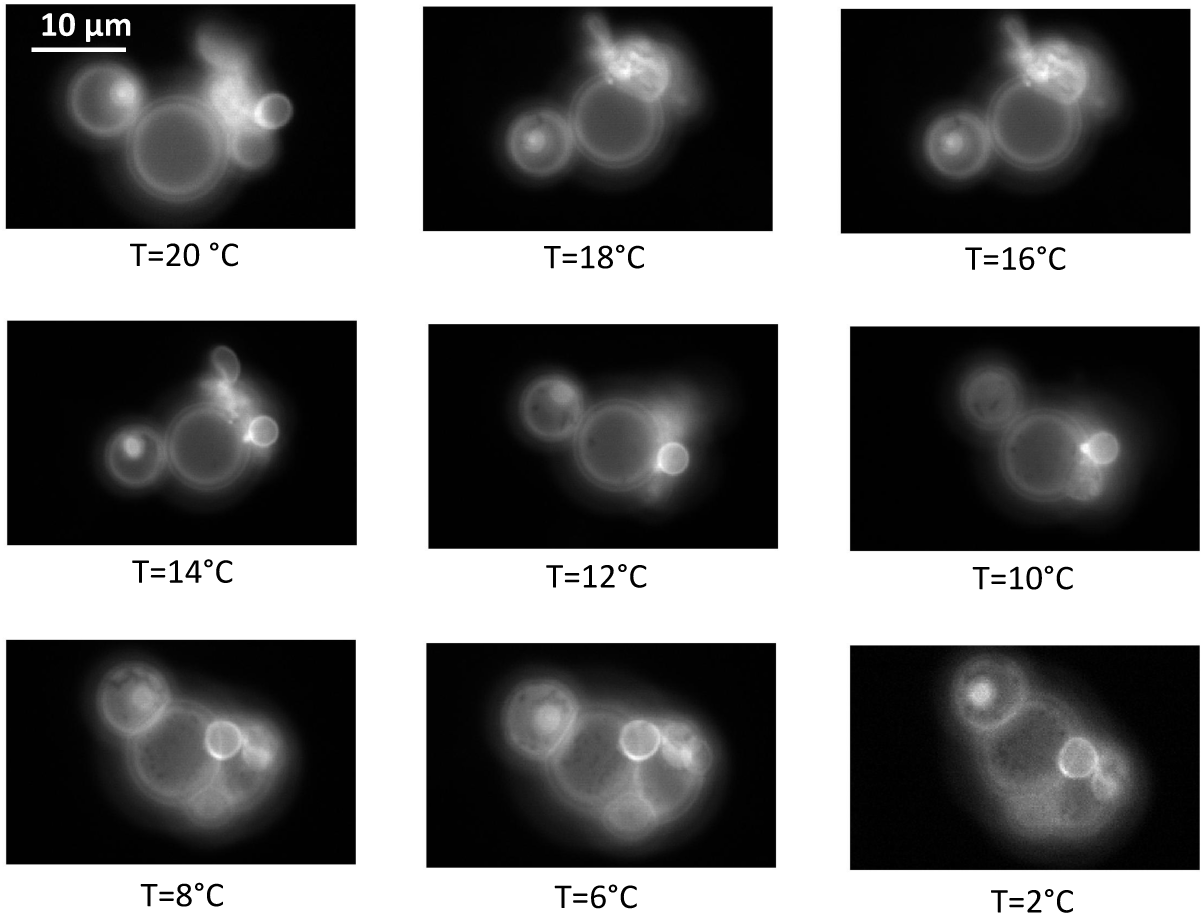
In the same way as the ‘parent’ GPMV, it has been checked that vesicles reformed from just the lipid fraction purified from GPMVs undergo phase separation. The images show phase separation (dark domains) appearing at 18 and 12°C.

